# Mechanotransduction events at the physiological site of touch detection

**DOI:** 10.1101/2022.10.23.513402

**Authors:** Luke H. Ziolkowski, Elena O. Gracheva, Sviatoslav N. Bagriantsev

## Abstract

Afferents of peripheral mechanoreceptors innervate the skin of vertebrates, where they detect physical touch via mechanically gated ion channels (mechanotransducers). While the afferent terminal is generally understood to be the primary site of mechanotransduction, the functional properties of mechanically activated (MA) ionic current generated by mechanotransducers at this location remain obscure. Here, we report patch-clamp recordings from the afferent terminal innervating Grandry (Meissner) corpuscles in the bill skin of a tactile specialist duck. We show that mechanical stimulation evokes MA current in the afferent with fast kinetics of activation and inactivation during the dynamic phases of the mechanical stimulus. These responses trigger rapidly adapting firing in the afferent detected at the terminal and in the afferent fiber outside of the corpuscle. Our findings elucidate the initial electrogenic events of touch detection in the mechanoreceptor nerve terminal.

## Introduction

In vertebrates, extrinsic touch is detected in the skin by cutaneous mechanoreceptors, somatosensory neurons of the peripheral nervous system. The afferent nerve fibers of these cells innervate the skin, where they form specialized ending structures which sense mechanical stimuli. Within the afferent terminals, mechanically-gated ion channels (mechanotransducers), such as Piezo2, detect touch and transform it into mechanically activated (MA) current (Handler and Ginty, 2021). Extracellular recordings of mechanoreceptor afferents have previously revealed voltage changes originating from the terminals in response to mechanical stimulation, but the intracellular dynamics of these signals are not understood (Loewenstein et al., 1958). As a result, direct evidence of mechanotransduction and MA current in the nerve endings of mechanoreceptors is lacking.

Studies of MA current and mechanotransducer biophysics have been limited to heterologous expression systems and dissociated somatosensory neurons *in vitro* (Coste et al., 2010; Lewis et al., 2017; Schneider et al., 2017; Zheng et al., 2019). Most notably, Piezo2, which mediates the detection of touch, displays fast-inactivating MA current in cultured cells and in dissociated neurons (Buchholtz et al., 2021; Chesler et al., 2016; Coste et al., 2010; Ranade et al., 2014; Wang et al., 2019). However, it is unclear whether electrophysiological responses of somas from dissociated neurons accurately reflect that of MA current in the afferent terminal, due to potential differences in membrane geometry, level of ion channel expression, intracellular factors, and cellular/tissue environment between the two (Richardson et al., 2022). To our knowledge, intracellular recordings of mechanoreceptor terminals have not been previously reported due to the technical difficulties of accessing the axonal endings with patch-clamp electrodes. Consequently, the functional characteristics of mechanotransduction at the normal physiological site of touch detection remain unknown.

To address this gap in knowledge, we acquired patch-clamp recordings from the afferent terminals of Grandry corpuscles in the bill skin of the tactile specialist Mallard duck (*Anas platyrhynchos domesticus*). The Grandry corpuscle is an avian tactile end-organ innervated by rapidly-adapting mechanoreceptors, which form thin terminals between Schwann cell-derived lamellar cells (Nikolaev et al., 2020; Schneider et al., 2017). The Grandry corpuscle’s layered architecture, rapid adaptation, and sensitivity to transient touch make it structurally and functionally analogous to the mammalian Meissner corpuscle (Neubarth et al., 2020; Schwaller et al., 2021; Ziolkowski et al., 2022). Compared to mammals, the high density of corpuscles in the bill of tactile-foraging waterfowl enables persistent electrophysiological investigation of the afferent terminals in these end-organs, the results of which we report here.

## Results and Discussion

We acquired patch-clamp recordings from the afferent terminal within the Grandry corpuscle using an *ex vivo* bill-skin preparation from late-stage duck embryos (Fig. 1A). Mechanical stimulation of the voltage-clamped afferent terminal revealed fast-inactivating MA current only in response to the dynamic onset (ON) and offset (OFF) phases of the stimulus (Fig. 1B). In current-clamp, both indentation with a probe (Fig. 1C) and current injection (Fig. 1D) caused depolarization of the membrane voltage, which initiated action potentials (APs) in the terminal during both phases. In three corpuscles in which the afferent terminal was patched, simultaneous single-fiber nerve recordings were also established using a section of the same afferent outside of the corpuscle (Fig 1A,E). In these cases, propagating action potentials from the afferent terminal were recorded in the afferent fiber with a one-to-one correlation to APs in the terminal (Fig. 1B-D, bottom). When comparing the responses during the ON and OFF phases, we detected a difference between the rates of current inactivation (Fig. 1F), but not the rates of activation, current-indentation relationship, or AP threshold (Fig. 1E-I). The inactivation rate of MA current in the ON phase (τ = 8.95 ± 1.82 ms) in the terminal is notably similar to the inactivation rate of fast-inactivating MA current measured from the somas of murine and duck mechanoreceptors *in vitro* (Coste et al., 2010; Schneider et al., 2017; Viatchenko-Karpinski and Gu, 2016). Interestingly, the MA current seen during the OFF phase is a unique response not reported in dissociated neurons or expression systems, even though the OFF response is typical of rapidly adapting mechanoreceptors in *ex vivo* single-fiber recordings. The fast inactivation rate of the OFF response compared to the ON response implies a distinct or modified mechanism of mechanotransduction. This could potentially be dependent on the cellular structure or function of lamellar cells in the corpuscle.

**Figure 1.**
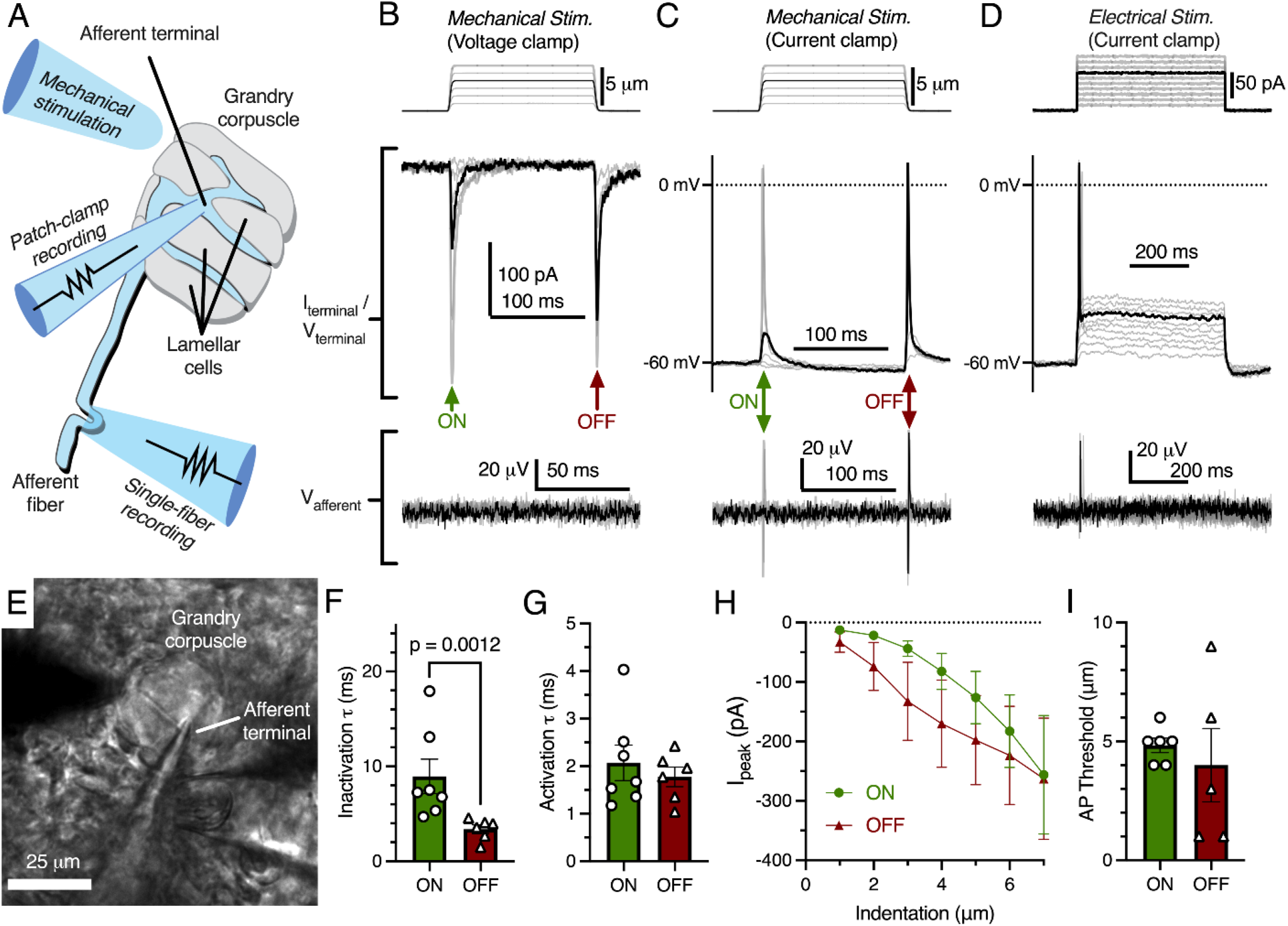
Mechanotransduction in the terminal afferent of Grandry corpuscle. **(A)** Illustrated representation of the experimental setup. **(B)** The mechanical step stimulus applied with a glass probe (top), representative MA current responses in the terminal while voltage-clamped at -60 mV (middle), and simultaneous extracellular voltage signal from the connected afferent (bottom). **(C)** The mechanical stimulus (top), voltage responses and APs in the terminal in current-clamp (middle), and APs measured further along the afferent (bottom). **(D)** The current injection stimulus (top), voltage responses and action potentials in the terminal in current-clamp (middle), and APs measured in the afferent (bottom). **(E)** Example bright-field image of the experimental setup. **(F)** Quantification of the kinetics of MA current inactivation, **(G)** activation, **(H)** peak MA current-indentation relationship (n = 7/6 for ON/OFF, respectively), and **(I)** AP threshold measured in the dynamic onset phase the stimulus (ON) and the dynamic offset phase of the stimulus (OFF). Only the difference in inactivation τ between the ON and OFF phase was statistically significant (p < 0.05). Statistics: Mann-Whitney U test (F,G,I) or two-way ANOVA (H). Symbols indicate data from individual cells. Data in F-I were obtained from at least 3 independent skin preparations, and shown as mean ± SEM.

As expected, the addition of tetrodotoxin (TTX) to the bill-skin preparation blocked APs and voltage-gated sodium current in the afferent terminal (Fig. 2A-D). In some voltage clamp experiments, mechanical stimulation resulted in large (>1000 pA) depolarizing currents (Fig. 2C) which did not follow the expected current-indentation relationship (Fig. 1H). These currents were blocked by TTX and therefore were voltage-gated sodium currents resulting from brief loss of voltage clamp, likely due to the complex geometry of the afferent.

**Figure 2.**
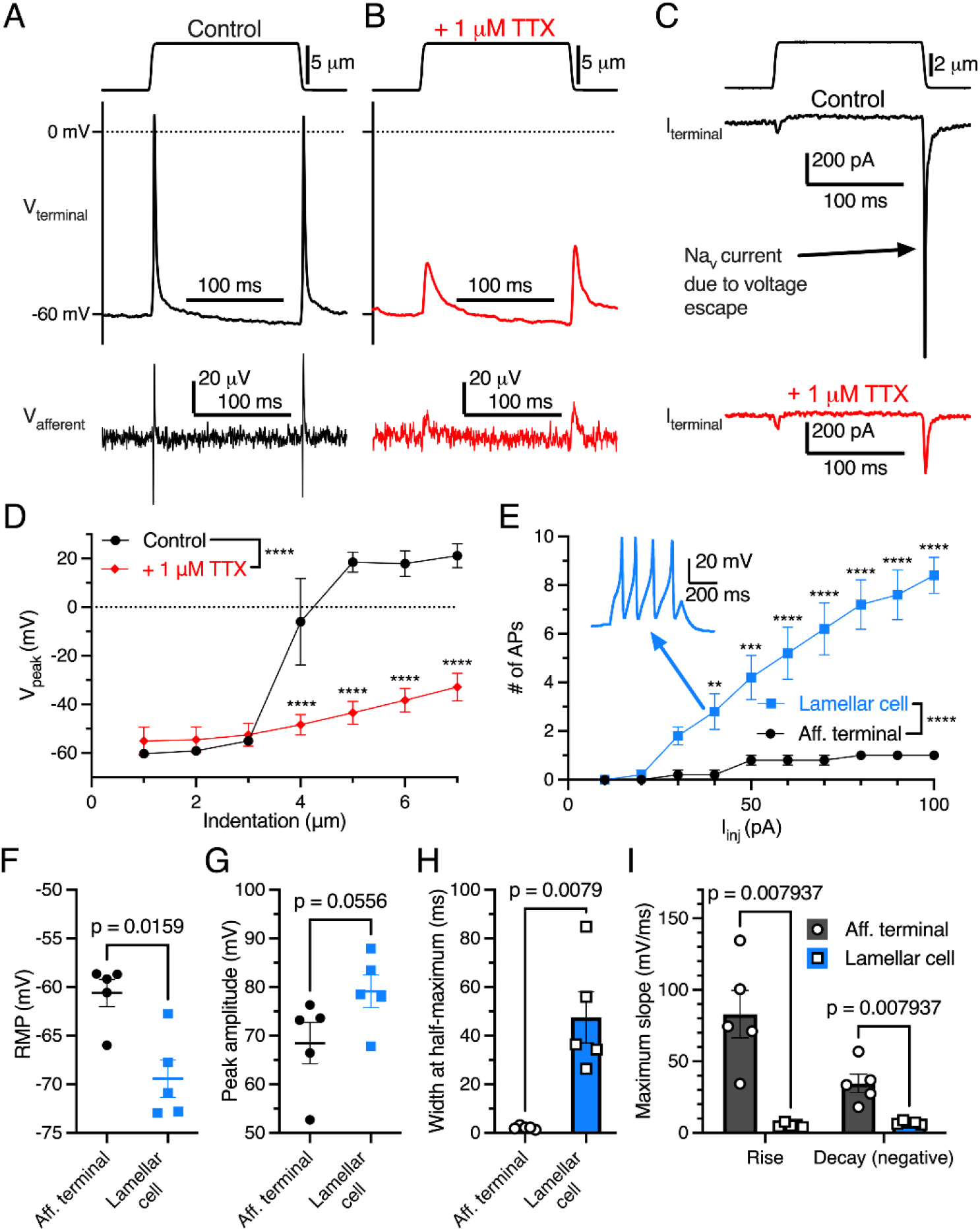
Electrogenic events in mechanoreceptor terminal and lamellar cells are carried out by different mechanisms. **(A)** A suprathreshold mechanical stimulus (top), APs in the terminal (middle), and propagated APs from the connected afferent (bottom). **(B)** A suprathreshold mechanical stimulus applied in 1 µM TTX (top), AP-absent voltage responses in the terminal in current-clamp (middle), and extracellular receptor potentials in the afferent (bottom). **(C)** A suprathreshold mechanical stimulus (top), current responses in the terminal while voltage-clamped at -60 mV without 1 µM TTX (middle), and with 1 µM TTX (bottom). **(D)** Voltage-indentation relationship in the absence or presence of 1 µM TTX (n = 5 for both groups). **(E)** The number of APs from increasing current injections in lamellar cells and afferent terminals (n = 5 for both groups). **(F)** Resting membrane potential (RMP), **(G)** peak AP amplitude, **(H)** AP width at the half-maximum, and **(I)** the maximum slope of the AP rise or decay in the afferent terminal versus lamellar cells of the corpuscle. The AP-current injection relationship, resting membrane potential, width at half-maximum, max rise slope, and max decay slope were significantly different between terminal afferent and lamellar cells (p < 0.05). Statistics: Mann-Whitney U test (F-I) or two-way ANOVA with Holm-Sidak post-hoc test (D,E). **P=0.0084, ***P=0.0004, ****P<0.0001. Symbols indicate data from individual cells. Data in D-I were obtained from at least 3 independent skin preparations, and shown as mean ± SEM.

Importantly, APs in the afferent terminal are physiologically distinct from APs fired by Grandry lamellar cells (Fig. 2E-I). Lamellar cell APs are mediated by voltage-gated calcium channels, which are insensitive to TTX (Nikolaev et al., 2020). Lamellar cells fire multiple APs in response to large current injections, whereas the afferent terminal fires a maximum of one AP during the same stimuli (Fig 2E). Additionally, there were significant differences in resting membrane potential, AP width at half-maximum, and maximum slope of rise and decay between the two cell types. These results, along with the single-fiber afferent voltage data which mirrors the terminal voltage (Fig 1B-D), demonstrate that the recordings acquired here are unequivocally from the afferent terminal within the corpuscle.

Here, we have shown that mechanical stimulation evokes MA current in the afferent terminal which initiates propagating APs. Critically, MA current in the terminal has properties closely resembling those observed in dissociated neuron somas. This ultimately confirms the validity of using *in vitro* models to study mechanotransducers. At the same time, an important aspect of the afferent terminal response *in situ* is absent from cultured cells: the MA current in the OFF phase. Further studies of rapidly adapting corpuscles and other mechanoreceptor endings will be required to understand the mechanism underlying the OFF response. Together, these findings reveal fundamental characteristics of mechanotransduction at the physiological site of touch detection in mechanosensory neurons.

## Supplemental Figure Legends

**Figure 1 – Source Data 1. Original data for**

**Figure 1F-I Figure 2 – Source Data 2. Original data for Figure 2D-I**

## Materials and Methods

### *Ex vivo* bill-skin preparation

Experiments with duck embryos (*Anas platyrhynchos domesticus*) were approved by and performed in accordance with guidelines of the Institutional Animal Case and Use Committee of Yale University, protocol 11526. Bill preparation was slightly modified from previously published methods (Nikolaev et al., 2020). Intact skin was carefully removed from the bill of duck embryos (aged embryonic day 25 to 27, sex not determined) using a sharp scalpel tip in ice-cold L-15 media. The bill-skin was placed upside-down (epidermis on bottom) in the recording chamber under a slice anchor. Corpuscles and afferents in the dermis were visualized on an Olympus BX51WI upright microscope with an ORCA-Flash 4.0 LT camera (Hamamatsu). At room temperature (22-23°C), the bill-skin preparation was treated for 5 minutes with 2 mg/mL collagenase P (Roche) in Krebs solution containing (in mM) 117 NaCl, 3.5 KCl, 2.5 CaCl_2_, 1.2 MgCl_2_, 1.2 NaH_2_PO_4_, 25 NaHCO_3_, and 11 glucose, saturated with 95% O_2_ and 5% CO_2_ (pH = 7.3-7.4), then washed with fresh Krebs solution.

### Patch-clamp electrophysiology

Recordings were at room temperature using a MultiClamp 700B amplifier, Digidata 1550A digitizer, and pClamp 10 software (Molecular Devices). Standard-wall, 1.5 mm diameter borosilicate pipettes with tip resistances of 2-5 MΩ were pulled using a P-1000 micropipette puller (Sutter Instruments). Pipettes were filled intracellular solution containing (in mM) 135 K-gluconate, 5 KCl, 0.5 CaCl_2_, 2 MgCl_2_, 5 EGTA, 5 HEPES, 5 Na_2_ATP, and 0.5 Na_2_GTP (pH 7.3 with KOH). All experiments were performed in Krebs solution at room temperature. Data were sampled at 20 kHz and low-pass filtered at 2 kHz. Terminals were recorded in whole-cell mode and were held at -60 mV during voltage-clamp experiments. Resting membrane potential was measured in current-clamp mode shortly after breaking in. In both voltage- and current-clamp, mechanical stimuli were applied to a single corpuscle using a blunt glass probe (2 to 10 µm tip diameter) mounted on a piezoelectric-driven actuator (Physik Instrumente GmbH. A mechanical step stimulus was applied to corpuscles starting at 1 µm and increasing by 1 µm after each indentation. The static plateau of the step stimulus lasted 150 ms, while the ramp had a duration of 3 ms for both the ON and OFF phases. For both phases in each terminal, the inactivation rate (τ) of the MA current was calculated by fitting the equation I = I_0_*exp^(− t/τ) to the decaying portion of the largest three MA current responses, and averaging those τ values. The activation τ was calculated similarly using the rise portion of the response. The threshold was measured in current-clamp as the smallest indentation which elicited an AP. In current-clamp, depolarizing current steps (from 10 to 100 pA in 10 pA increments) were applied to elicit APs in the afferent terminal and lamellar cells. The first AP in these recordings was used to calculate the peak amplitude, width at half-maximum and maximum slope of rise and decay for the terminal versus lamellar cells. Experiments were not corrected for liquid-junction potential.

### Single-fiber recording

Recordings from single afferent fibers of corpuscles were acquired simultaneously with patch-clamp recordings for three corpuscles, using the second channel of the MultiClamp 700B amplifier. Single-fiber recording pipettes were created pulled from thin-wall, 1.5 mm diameter borosilicate glass capillaries using a P-1000 micropipette puller (Sutter Instruments) to create tip diameters of 5 to 30 µm, then filled with Krebs solution. Pipettes were placed on an electrode headstage connected to a High Speed Pressure Clamp (ALA Scientific Instruments). Light (1 to 20 mmHg) positive pressure was applied from the recording electrode to clear away tissue from a corpuscle-associated afferent. Negative pressure was then applied until a large section (∼5 µm) of the afferent was sucked into the pipette. Extracellular afferent voltage was recording in current-clamp mode, sampled at 20 kHz and low-pass filtered at 1 kHz.

### Data analysis

Data from afferent terminals and lamellar cells were acquired from separate, individual preparations from different animals. Data were analyzed and plotted in GraphPad Prism 9.4.1 (GraphPad Software, LLC) as individual data points or means ± SEM, unless otherwise indicated.

## Acknowledgements

We thank Dr. Yury Nikolaev for help with establishing the skin preparation, and members of the S.N.B. and E.O.G. laboratories for their contributions throughout the project. This study was partly funded by NSF grants 1923127, 2114084 (to S.N.B) and 1754286 (to E.O.G.), and NIH grants R01NS097547 and R01NS126277 (to S.N.B).

